# DeepES: Deep learning-based enzyme screening to identify orphan enzyme genes

**DOI:** 10.1101/2024.05.09.592857

**Authors:** Keisuke Hirota, Felix Salim, Takuji Yamada

**Affiliations:** School of Life Science and Technology, Tokyo Institute of Technology, Tokyo, Japan; Metagen, Inc., Yamagata, Japan; Metagen Therapeutics, Inc., Yamagata, Japan; digzyme, Inc., Tokyo, Japan

## Abstract

**Motivation:** Progress in sequencing technology has led to determination of large numbers of protein sequences, and large enzyme databases are now available. Although many computational tools for enzyme annotation were developed, sequence information is unavailable for many enzymes, known as orphan enzymes. These orphan enzymes hinder sequence similarity-based functional annotation, leading gaps in understanding the association between sequences and enzymatic reactions.

**Results:** Therefore, we developed DeepES, a deep learning-based tool for enzyme screening to identify orphan enzyme genes, focusing on biosynthetic gene clusters and reaction class. DeepES uses protein sequences as inputs and evaluates whether the input genes contain biosynthetic gene clusters of interest by integrating the outputs of the binary classifier for each reaction class. The validation results suggested that DeepES can capture functional similarity between protein sequences, and it can be implemented to explore orphan enzyme genes. By applying DeepES to 4744 metagenome-assembled genomes, we identified candidate genes for 236 orphan enzymes, including those involved in short-chain fatty acid production as a characteristic pathway in human gut bacteria.

**Availability and implementation:** DeepES is available at https://github.com/yamada-lab/DeepES. Model weights and the candidate genes are available at Zenodo (https://doi.org/10.5281/zenodo.11123900).

## 1. Introduction

Progress in sequencing technology has yielded large numbers of protein sequences, including enzyme sequences. Although large enzyme databases, such as Kyoto Encyclopedia of Genes and Genomes (KEGG) (Kanehisa *et al*. 2021) and the BRENDA (Chang *et al*. 2021), are available, many enzymes registered in these databases lack sequence information. Enzymes with confirmed activity for which associated protein sequence information is lacking are termed orphan enzymes (Lespinet and Labedan 2005; Pouliot and Karp 2007). Reactions involving orphan enzymes involved in 20.8% of the 11 914 metabolic reactions described in the KEGG database (KEGG FTP Release 27 March 2023). Furthermore, at least 22.4% of enzymes with Enzyme Commission (EC) numbers are orphans (Sorokina *et al*. 2014). Leveraging genome-context information is useful for identifying orphan enzyme genes (Yamada *et al*. 2012); however, a computational approach for this identification has not been developed.

Orphan enzymes cannot be detected using sequence similarity-based functional annotation approaches, such as BLAST (Altschul *et al*. 1990), because their sequences are unknown. This issue severely limits biological function studies based on molecular data and creates gaps between the enormous sequence data available and the understanding of biological processes (Yamada *et al*. 2012, Zhang and Li 2023). For instance, diverse metabolic processes performed by the human gut microbiota, such as short-chain fatty acid production, influence gut inflammation and cancer progression (Parada Venegas *et al*. 2019, Rebersek 2021, Liu *et al*. 2023); however, many of these reactions involve orphan enzymes (Shiroma *et al*. 2023). Therefore, an approach that assigns enzymatic activities to gene sequences without calculating sequence similarity is essential for understanding the role of orphan enzymes.

Functions can be predicted from a gene sequence independently of reference sequence databases. Recently, deep learning has enabled prediction of protein 3D structures (Jumper *et al*. 2021, Lin *et al*. 2023) and function classification from a protein sequence with high accuracy (Sureyya Rifaioglu *et al*. 2019, Cao and Shen 2021, Littmann *et al*. 2021, Sanderson *et al*. 2023). Most deep learning-based approaches focusing on enzymes, including DeepEC (Ryu, Kim and Lee 2019), CLEAN (Yu *et al*. 2023), EnzBert (Buton, Coste and Le Cunff 2023) and DeepECtransformer (Kim *et al*. 2023), predict enzymatic activity based solely on a protein sequence in a powerful and efficient manner. EC numbers have also been used as enzymatic activity classes. The EC number, the most well-known enzyme classification system, describes enzymatic reactions hierarchically using four-digit numbers. Although the EC number is useful, the specified EC number require complete description of the enzymatic activity. Therefore, it is incompatible with orphan enzymes, as the activities of these enzymes are often unclear. To overcome this limitation, we utilized Reaction Class (RClass), a classification system based on chemical transformation patterns for substrate and product pairs (Kotera *et al*. 2004, Muto *et al*. 2013). RClass can be used to classify incomplete reactions because classifications are defined from the fingerprint of a substrate-product pair, making this approach more appropriate than using EC numbers for orphan enzymes.

In this study, we developed DeepES, a deep learning-based framework focusing on sequential orphan enzymes using biosynthetic gene clusters (BGCs) for *in silico* orphan enzyme screening using RClass (Fig. 1). DeepES utilizes ESM-2, a pre-trained large language model that convert protein sequences to vector representations (Lin *et al*. 2023). We show that DeepES can identify candidate genes for orphan enzymes. DeepES consists of RClass binary classifiers, which predict whether the input genes correspond to each RClass, and the outputs of each model are integrated to evaluate whether the input genes contain the BGC, including an orphan enzyme of interest. Subsequently, we focused on human gut bacteria and explored candidate genes from 4744 metagenome-assembled genomes (MAGs) for 236 orphan enzymes, including enzymatic reactions for short-chain fatty acid production, as characteristic pathways of human gut bacteria.

**Figure 1.**
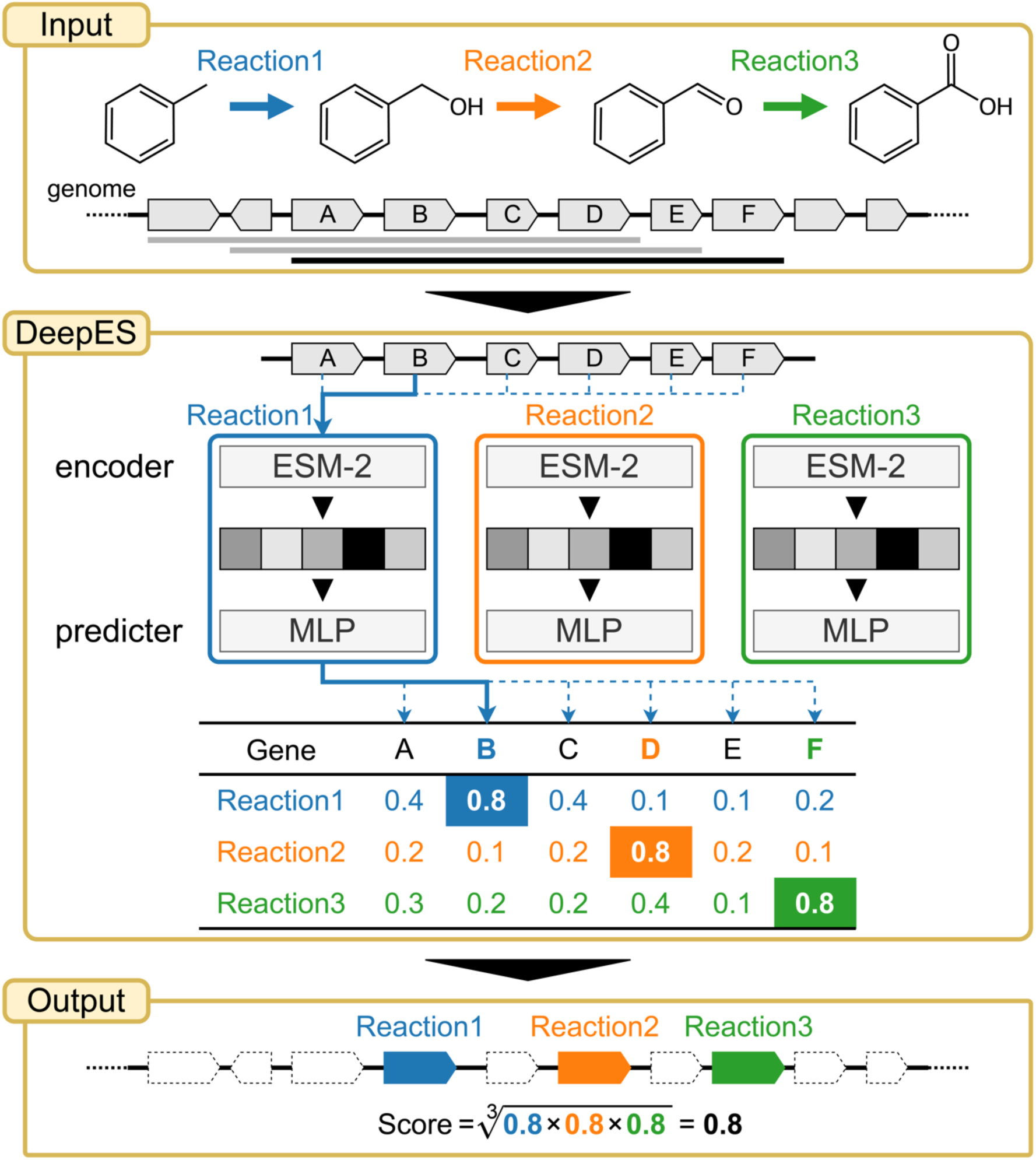
DeepES overview. The flow chart of exploring candidate genes for sequential enzymes using DeepES (our proposed method) consists of a three-step process: (1) DeepES retrieves protein sequences of successive genes in the genome from the input gene list, (2) infers independently the probability that each gene corresponds to each enzyme using the RClass classifiers for the enzyme reactions of interest and (3) evaluates whether the sequential genes are likely to encode the enzymes for given successive chemical reactions based on a geometric mean of predicted probabilities. MLP, multilayer perceptron.

## 2. Materials and Methods

### 2.1. Datasets

We used prokaryotic gene sequences and RClass data retrieved from the KEGG database (KEGG FTP Release 27 March 2023) for the training, validation, and testing of machine learning (ML) models. KEGG Orthology (KO) is a dataset containing the corresponding enzymatic reactions from a gene to a protein orthology. Therefore, a dataset for ML models was constructed by pairing gene sequences and RClass using KO; the data included 4 413 823 cases. Protein sequences of more than 4000 amino acids or those containing "J" (leucine or isoleucine) were removed from the datasets because of the computational resources and specifications of ESM-2. The final dataset consisted of 4 413 788 prokaryotic gene protein sequences and RClass pairs split into 1 896 609 training data points and 2 517 179 test data points as of 15 May 2017.

### 2.2. RClass

RClass is a fingerprint-based reaction classification schema and KEGG RCLASS database entry (Kotera *et al*. 2004, Muto *et al*. 2013). The KEGG RCLASS database contains all chemical transformation patterns associated with all reactions in the KEGG REACTION database. Each RClass represents the chemical transformation pattern of a substrate-product pair, and one reaction is often annotated with multiple RClasses. RClass allows a more abstract and automatic classification than the EC number does, and thus, can be applied to a wider range of enzymatic reactions.

### 2.3. ML model development

RClass prediction is a multi-label classification task because several RClasses can be annotated for an enzyme reaction. RClass is also an imbalanced classification, with only 475 of the 1454 RClasses accounting for 80% of the data. Therefore, the ML models constituting DeepES were implemented as binary classifiers for each RClass. Each ML model consists of ESM-2 (Lin *et al*. 2023), a multilayer perceptron, and a simple feedforward fully connected neural network; that is, the first half embeds the input gene sequence into a vector representation, and the second half predicts the probability that the input gene corresponds to the RClass of interest based on its embedding vector. We used ESM-2 with the specific version esm2_t33_650M_UR50D and froze its parameters. We also adopted a weighted loss function based on the class weight to address the class imbalance of RClass. If the enzyme reaction had multiple RClasses, the geometric mean of the model output corresponding to each RClass was calculated as the predicted probability, which indicated whether a given protein sequence encoded the enzyme.

To determine the optimal hyperparameters, we performed a grid search with 10-fold cross-validation on the training data and compared the area under the precision-recall curve (AUPRC). The hyperparameters included the number of epochs, hidden layer size, learning rate, and dropout rate (Table 1). We performed hyperparameter optimization of the multilayer perceptron for 100 RClass with over 1000 data points because the training data contained approximately 1500 RClasses.

**Table 1.**
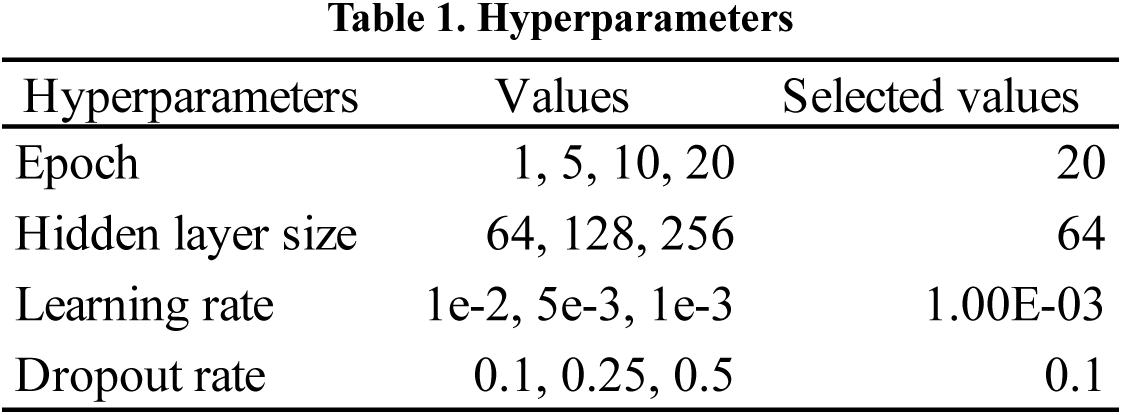
Hyperparameters.

### 2.4. Validation using BLAST

We examined whether the developed ML models could predict enzymatic functions that are difficult to annotate using BLAST. The hyperparameters were optimized as described in 2.3.

First, we created a small non-redundant validation dataset via sequence homology clustering of the KEGG test dataset using MMseqs2 (Steinegger and Söding 2017) with an identity threshold of 0.3 and the other parameters set to default values. We then repeated the steps of (1) constructing the training and test datasets, (2) training the ML models, and (3) evaluating ML model performance in a leave-one-out cross-validation manner using the validation dataset (Supplementary Fig. S1). Specifically, we chose one sequence (query sequence) and performed a homology search for the remainder of the dataset using BLAST (default) with the query sequence. We constructed a training dataset that excluded the hit sequences, and a test dataset by combining the query sequence data and negative data, which were 20 randomly selected non-enzyme genes from *Escherichia coli* K-12 MG1655. We trained the ML models and then evaluated their performance using the datasets. These procedures enabled us to calculate precision and evaluate whether the ML model output high predicted probabilities for positive examples and low predicted probabilities for negative examples. We also evaluated the reliability of the results of binary classification with a threshold for the predicted probability.

### 2.5. Biosynthetic gene cluster detection

In bacterial, fungal, and plant genomes, groups of genes encoding enzymes, transporters, and regulators associated with a series of chemical reactions are often located close to each other in the genome; these gene are defined as BGCs, and are important for understanding metabolism in cellular systems (Medema *et al*. 2015, Kautsar *et al*. 2021). Furthermore, leveraging a combination of metabolic pathway adjacency and genomic neighborhood information can be used to obtain candidate genes of orphan enzyme genes (Yamada *et al*. 2012). Based on these findings, we developed DeepES, a deep learning-based framework focusing on BGCs for *in silico* orphan enzyme screening (Fig. 1). DeepES assumes that orphan enzyme genes constitute BGCs and aims to efficiently detect candidate genes. This framework takes a list of protein sequences of contiguous genes as input and repeats their evaluation using RClass classifiers corresponding to a series of enzyme reactions of interest. Subsequently, the set of contiguous genes slides the frame one by one from one end of the input gene list to the other and outputs a score representing the predicted probability that the BGC of interest falls within the range for each frame. The score for genes in the fixed scope of the input gene set was calculated by acquiring the highest predicted probability for each enzyme without gene overlap and calculating their geometric mean. For simplicity, the length of the contiguous reactions was 3, and the range of continuous genes evaluated was 10.

To validate the above BGC-based approach, 70 BGCs from the KEGG MODULE (KEGG FTP Release 27 March 2023) were used as pseudo-orphan enzymes. By preliminarily removing the KOs comprising these BCGs during training, we tested whether the proposed method could detect the candidate genes for pseudo-orphan enzymes. The 70 BGCs comprise a subset of the metabolic pathways in the KEGG MODULE that are detectable in this DeepES setting; that is, a group of 10 successive genes in the genome containing three enzymes corresponding to three sequential enzyme reactions in the metabolic pathway are present in the test data (Supplementary Table S1). For simplicity, we excluded BGCs showing non one-to-one correspondence between enzyme genes and enzyme reactions.

### 2.6. Application to real orphan enzymes

We applied DeepES to actual orphan enzymes to identify genes encoding an orphan enzyme. We specifically searched for candidate genes from 4744 representative MAGs of human gut bacteria obtained from MGnify (Unified Human Gastrointestinal Genome v2.0.1) (Richardson *et al*. 2023) for 332 orphan enzymes in EnteroPathway, the metabolic pathway database of the human gut microbiota (Shiroma *et al*. 2023). A total of 1235 orphan enzymes was collected as enzyme reactions that were not associated with sequence information based on the correspondence of each entry with KEGG or UniProt provided by EnteroPathway, and 332 enzymes were selected as DeepES-applicable; that is, 332 orphan enzymes were included in three consecutive enzyme reactions with RClass (Supplementary Table S2). We annotated RClass using E-zyme2 (Moriya *et al*. 2016) for enzymatic reactions in EnteroPathway when the RClass was not available on KEGG. Proteins with over 4000 residues or containing "J" (leucine or isoleucine) residue were excluded from the dataset because of limitations related to computational resources and specifications of ESM-2 when training and using the RClass classifiers.

## 3. Results

### 3.1. ML model development

After a grid search with 10-fold cross-validation, we selected the best hyperparameters by measuring the average AUPRC (Table 1, Supplementary Table S3) and evaluated the performance of the ML models using the test dataset. For the best hyperparameters, the RClass binary classifiers exhibited a high AUC and AUPRC on the test dataset (Fig. 2a).

**Figure 2.**
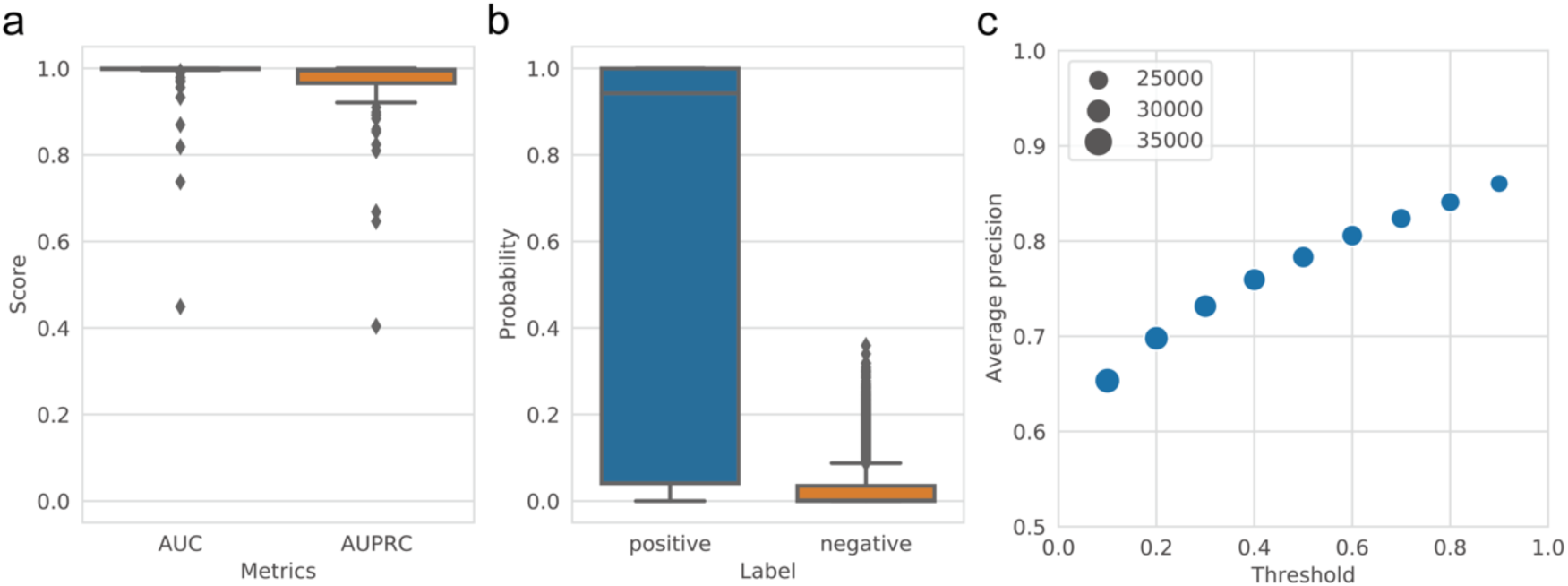
Performance of RClass binary classifier. (a) Predictive performance of the RClass binary classifiers on the test data (100 RClasses) evaluated by area under the receiver operating characteristic curve (AUC) and area under the precision-recall curve (AUPRC). (b) Distribution of predicted probabilities output by RClass binary classifiers in validation using BLAST. “Positive" and "negative" are output for query sequences and negative data, respectively. (c) Relationship of precision and threshold to predicted probability in validation using BLAST. The marker size represents the number of trials where precision could be calculated in leave-one-out cross-validation.

We next evaluated the predictive power of the ML models compared with that of BLAST by filtering the training dataset using BLAST as a leave-one-out cross-validation method. Although similar sequences in the test data were removed from the training dataset, the ML models provided high probabilities of predicting the correct RClass. The ML models showed high precision and appeared to clearly distinguish between positive and negative examples (Fig. 2b). Increasing the threshold for the predicted probability to 0.9 led to an average precision of 0.86, supporting the reliability of the upper end of the classifier’s prediction results (Fig. 2c).

### 3.2. DeepES performance test

We treated known enzymes as orphan enzymes by removing the corresponding genes from the training dataset and tested whether DeepES could detect candidate genes for these enzymes. If the KO of an enzyme was removed from the training data and treated as an orphan enzyme, the positive examples for the RClass, which is the label for that enzyme, were sometimes missing from the training data, and the classifier could not be built. When the enzyme corresponding to the middle of three consecutive enzyme reactions was treated as an orphan enzyme, 36 of 70 were testable (hereafter referred to as “one orphan”); when all three were treated as orphan enzymes, 19 of 70 were testable (hereafter referred to as “all orphans”) (Supplementary Table S1).

The results of applying DeepES to 36 BGCs (one orphan) and 19 BGCs (all orphans) showed that DeepES was able to detect the correct genes for the masked enzymes except for some BGCs (Supplementary Fig. S2, Supplementary Table S1). We selected the BGC with the largest number of positive cases per module to prevent duplication when evaluating enzymes and focused on those results (Fig. 3). Precision improved as the threshold for the DeepES output was increased; when the threshold was set to 0.99, the precision reached 0.690. Thus, approximately 70% of the scores above 0.99 in validation completely detected the three orphan enzyme genes, suggesting that the DeepES predictions with high scores are reliable. When all three genes selected by DeepES were hypothetical proteins, we excluded them from precision calculations because of evaluation difficulties. Furthermore, precision slightly improved when cases in which DeepES identified hypothetical proteins were considered. This result indicates that DeepES made an error and that a new gene for the enzyme could have been discovered.

**Figure 3.**
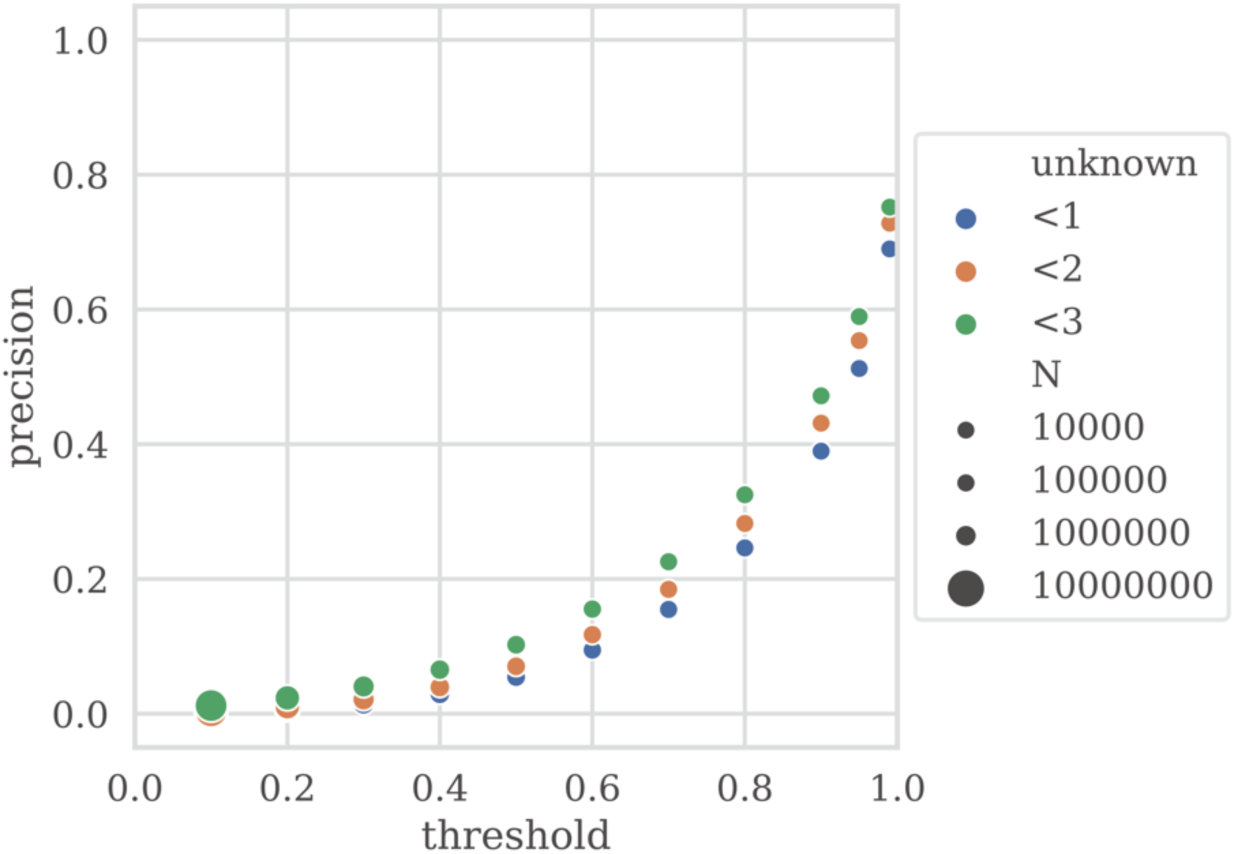
Validating DeepES on biosynthetic gene clusters (BGCs) with all orphans. Precision was calculated by summarizing the results of the DeepES predictions when the BGCs were selected from the 19 BGCs (all orphans) such that there was no overlap with the KEGG module, and the number of correct cases was maximized. The marker color indicates the number of hypothetical proteins that can be included in the genes hit by DeepES, and the marker size represents the number of outputs predicted as positive.

### 3.3. Application to actual orphan enzymes

We focused on human intestinal bacteria and explored candidate genes for orphan enzymes using DeepES. We applied DeepES to 4744 representative MAGs from MGnify (Unified Human Gastrointestinal Genome v2.0.1) (Richardson *et al*. 2023) to explore candidate genes for 332 orphan enzymes to which DeepES is applicable and are associated with gut bacteria-specific metabolism in the EnteroPathway (Shiroma *et al*. 2023). DeepES identified candidate genes for orphan enzymes of interest, as in the BGC detection test. DeepES was used to annotate 3 102 987 genes with 236 orphan enzymes involved in metabolic pathways unique to human gut bacteria, including short-chain fatty acid production (Table 2, Supplementary Table. S2). The BGC-based approach of DeepEC enabled us to narrow down candidate genes for orphan enzymes compared to evaluating a single gene and a single reaction (Fig. 4).

**Figure 4.**
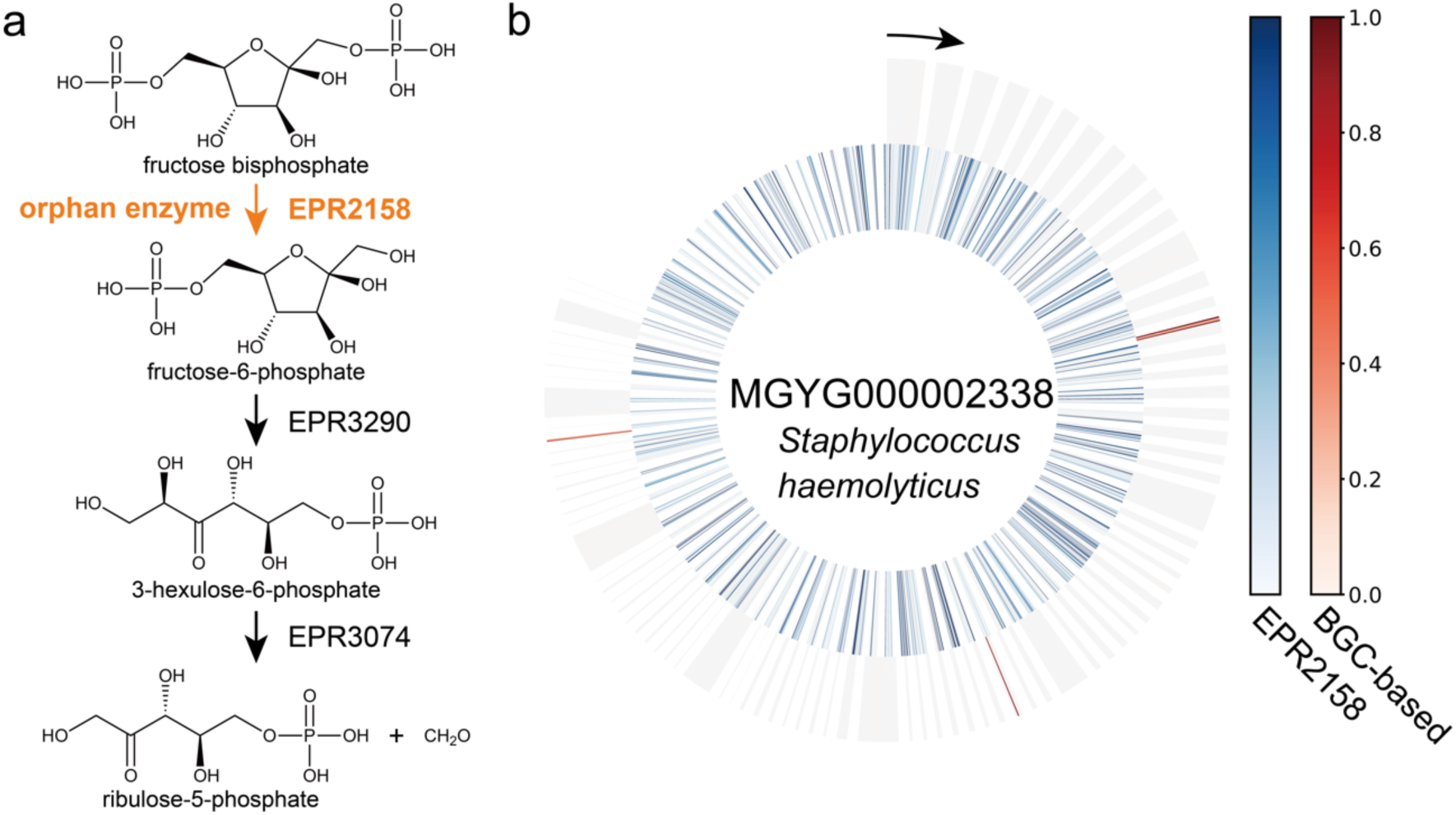
Example of orphan enzyme gene exploration using DeepES. (a) Example of part of a metabolic pathway involving enzymatic reactions by orphan enzymes. This part of energy metabolism by methane producers (EPM0948) and the first reaction (EPR2158) is performed by orphan enzymes. (b) Example of the result of applying DeepES to metagenome-assembled genomes (MAGs). The results for the MAG (MGYG000002338) with the highest score for the sequential enzyme reactions shown in (a) are illustrated. The inner circle (blue heatmap) represents the results of evaluating the orphan enzyme using only one RClass classifier, and the outer circle (red heatmap) represents the results of evaluating the series of enzyme reactions using DeepES. The window size of DeepES is 10.

**Table 2.**
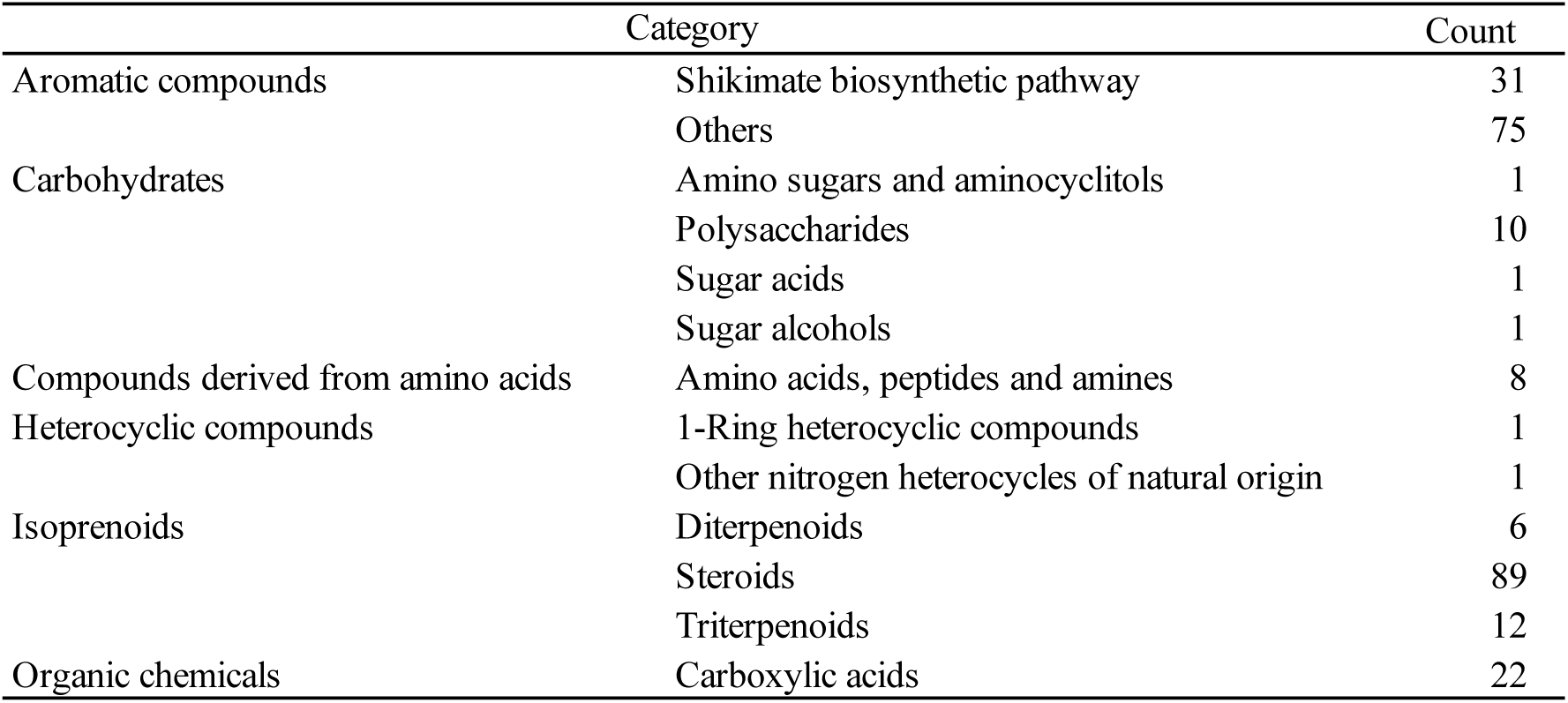
Categories of pathways involving orphan enzymes with candidate genes.

## 4. Discussions

We developed DeepES, a deep learning-based framework for enzyme screening focusing on BGCs and the RClass to identify orphan enzyme genes and demonstrated that the BGC-based approach may be useful for searching for orphan enzyme genes (Table 3). Evaluation of continuous genes inspired by BGCs is helpful for precise gene screening (Fig. 4). Unlike previous approaches, DeepES can screen orphan enzyme genes. Furthermore, the validation results using BLAST indicated that the ML model constituting DeepES is more sensitive than BLAST to functional similarity between sequences.

**Table 3.**
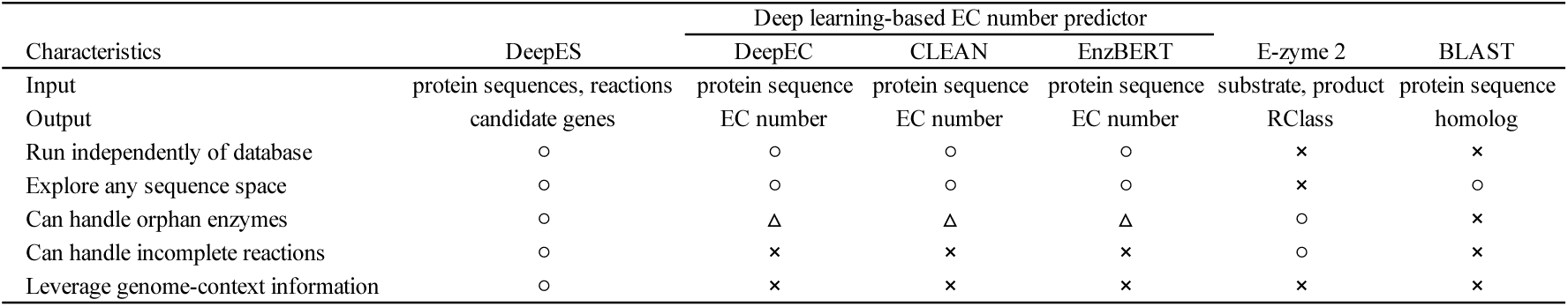
Comparison of DeepES with existing tools.

DeepES still has several limitations, including its representation of enzymatic activities and experimental validation. The use of RClass as a label for enzymatic activities allows ML models to predict enzymes without sufficient EC numbers, including orphan enzymes, whereas the class imbalance in the labels remains similar to that when using EC numbers. Although RClass is a fingerprint-based, high-quality reaction classification and can handle a wider range of enzyme reactions than can EC numbers, not all enzymatic reactions are annotated with RClass because RClass definition partially involves manual effort. Furthermore, RClass allows an approximate classification, which often leads to the assignment of the same RClass set to different enzyme reactions. Thus, the use of RClass can result in excessive candidate genes being hit, regardless of the performance of the RClass classifiers. However, no chemical reaction classification method is more automatic and more broadly applicable than, and of comparable quality to, RClass.

Many of the 236 orphan enzymes with candidate genes are involved in metabolic reactions associated with aromatic compounds or isoprenoids (Table 2). Some orphan enzyme reactions involved multiple pathways with different functions, such as EPR0399, which involved two functionally different pathways: organic chemicals (EPM0518) and aromatic compounds (EPM0665). Additionally, DeepES identified candidate genes for orphan enzymes involved in short-chain fatty acid production. Identifying genes corresponding to these enzymatic reactions may have many implications for studies of endogenous compounds, host-microbiota interactions (Rebersek 2021), and host metabolism (Gómez-Juaristi *et al*. 2018). MGYG000002338 (*Staphylococcus haemolyticus* strain:864-1) scored highest in EPR2158, the enzymatic reaction involved in short-chain fatty acid production (Fig.4). This predicted result may be promising, because some strains of Staphylococcus haemolyticus have been reported to produce an acid from fructose (Takeuchi *et al*. 2005). The prediction results for the orphan enzyme genes require experimental validation.

In conclusion, we developed DeepES, a novel tool to identify orphan enzyme genes by leveraging BGCs information and fingerprint-based enzyme classification. DeepES identified candidate genes for 236 orphan enzymes associated with human gut bacteria, indicating the potential of DeepES to bridge the gap between sequence data and biological understanding.

