## Supplementary material for "DeepES: Deep learning-based enzyme screening to identify orphan enzyme genes": Suppelementary_materials.pdf

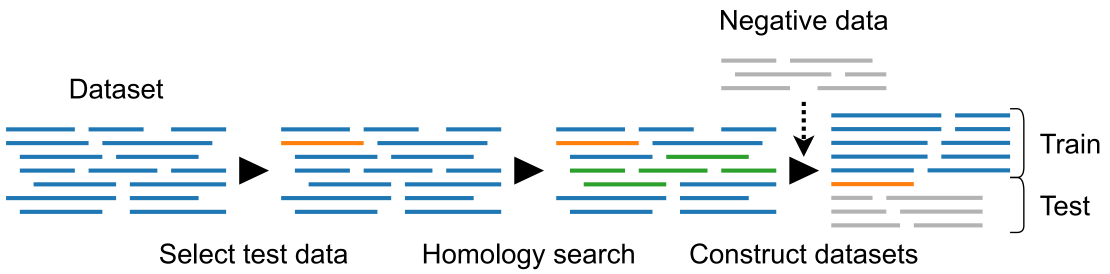


**Figure S1 Workflow to construct validation datasets using BLAST.** This process is repeated in leave-one-out cross-validation.

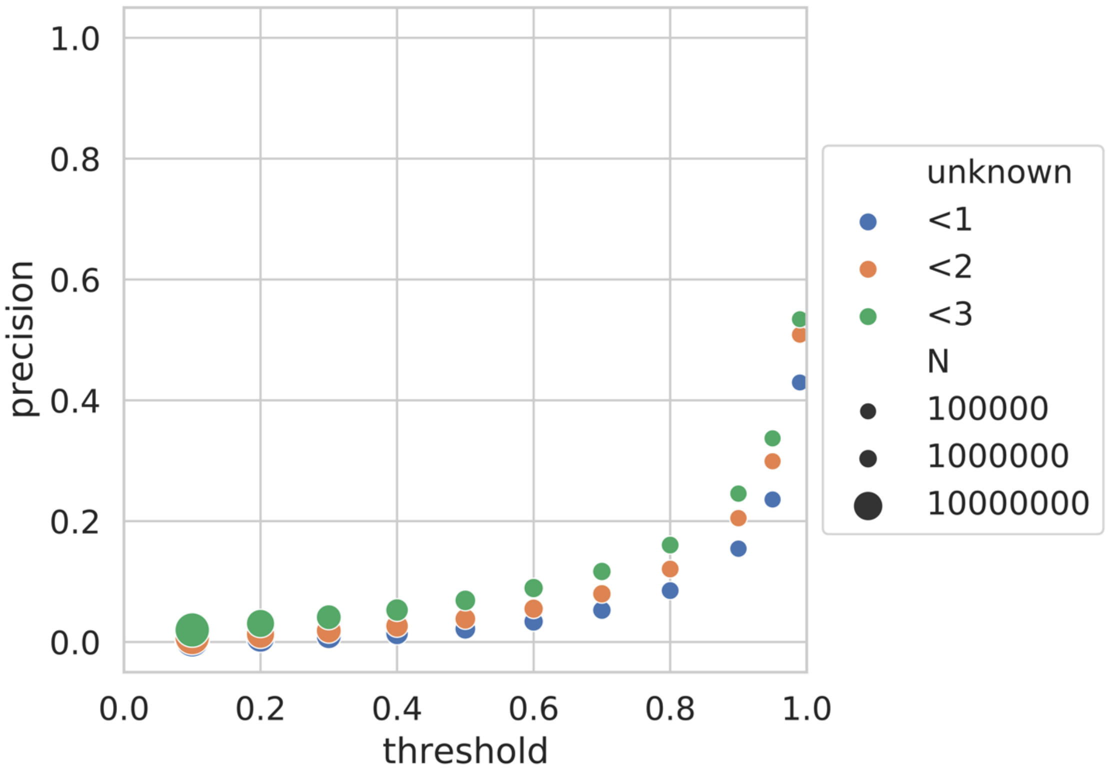


**Figure S2 Validating DeepES on BGCs with one orphan.** The precision was calculated by summarizing the results of the DeepES predictions for the 36 BGCs (one orphan). The marker color indicates how many hypothetical proteins can be included in the genes hit by DeepES, and the marker size represents the number of outputs predicted as positive.
